# P-DOpE probes reveal local amplification of phasic noradrenergic release in the hippocampus

**DOI:** 10.64898/2026.06.29.734897

**Authors:** Hengji Huang, Karyn Schy, Yue Liu, Alexandra Sommer, Jongwoon Kim, Brandon Jackson, Xintong Yang, Joanna Mattis, Earl T. Gilbert, Lijin Kevin Wang, Chelsea Buhler, Xiaoting Jia, Daniel Fine English, Sam McKenzie

**Affiliations:** Virginia Tech Bradley Department of Electrical and Computer Engineering, Blacksburg VA 24060; Virginia Tech School of Neuroscience, Blacksburg VA 24060; UNM Department of Neurosciences, UNM HSC, Albuquerque NM 87131; Department of Neurology and Michigan Neuroscience Institute, University of Michigan, Ann Arbor, Michigan, USA

## Abstract

Fiber photometry (FP) has become a tool of choice for in vivo monitoring of genetically encoded biosensors. The ability to record and optogenetically manipulate circuits through the same fiber stub is powerful, but limited, as biosensors typically do not sample membrane voltage, leaving the experimenter blind to the direct effects of opsin photoactivation. Here we developed the Photometry Device with Optogenetics and Electrophysiology (P-DOpE) probe, fabricated via a new convergence taper-break (CTB) method that integrates industry-standard silica optical waveguides with low-impedance metal electrodes that can be arranged in experimenter-defined configurations. We demonstrate that chronically implanted P-DOpE probes provide months-long recordings of local field potential, single unit recording, and fiber photometry, with parallel optogenetic circuit perturbation. Conducting fiber photometry with same-site optogenetic stimulation, we identified a robust fluorescence signal that scaled with network activity and survived biosensor antagonism. As this confound could not be eliminated with standard isosbestic controls, we propose a simple correction strategy. As a first application, we used the probe to test a proposed mechanism for focal modulation of noradrenergic signaling in and by cortical circuits receiving afferents from the locus coeruleus. We found that increasing spiking activity in CA1 amplifies noradrenergic signaling evoked by contextual arousal by ∼50%, but does not induce norepinephrine release in the absence of a phasic trigger – thus supporting the central prediction of the glutamate amplifies noradrenergic effects (GANE) hypothesis. The P-DOpE probe thus enables optogenetic manipulation and multimodal readout in a configurable low-cost, scalable, and robust format.

## Introduction

Communication in the brain relies on the interaction between rapid fluctuations of the membrane potential and the release of the hundreds of intra- and extracellular signaling molecules that directly and indirectly control transmembrane ion flux. While mature tools exist for monitoring electrical activity, measuring neuromodulatory dynamics in behaving animals has historically been limited to methods with poor temporal resolution, such as microdialysis, or low signal-to-noise, such as electrochemistry^1,2^. More recently, genetically encoded fluorescent indicators (GEFIs) have been developed that track substrate binding for classical and non-classical neurotransmitters, as well as intracellular signaling pathways^3-5^. The recent explosion of ligands for which GEFIs exist, and the simple and cost-effective nature of fiber photometry (FP), has made this a tool of choice for many neuroscience applications^6,7^. Alongside developments in measurement, a growing suite of optogenetic tools now provide unprecedented control of neuronal electrical activity^8^. Combining optogenetic manipulation and FP monitoring allows for precise analysis of how various cell populations and regions control extracellular and intracellular signaling. Despite rapid advances in bioengineering, there remains no standardized instrumentation for optogenetic perturbation that simultaneously captures immediate electrophysiological responses alongside concurrent non-electrical changes in the neurochemical milieu.

Attempts to integrate photometry, optogenetics, and electrophysiology into a single neural probe have typically used separate or bonded components^9,10^, which are difficult to scale for commercial production and often do not allow reliable co-localization of device elements. The thermal drawing process offers scalable fabrication of multimodal fiber-based probes^11-16^ but faces key challenges surrounding electrode count and material incompatibilities that hinder integration of high-performance materials like silica with metal electrodes. Our recent T-DOpE probe used this approach to integrate optogenetics and electrophysiology in a single fiber but did not provide photometry^17^.

To overcome these limitations, we introduce the P-DOpE probe based on a novel convergence taper-break (CTB) method that exploits the Marangoni effect to integrate dissimilar materials like silica, metal, and polymers within miniaturized fiber probes. We show that this scalable, low-cost approach enables probes with up to 28 electrodes and industry-standard silica waveguides. The electrodes consist of a variety of high-melting point metal or metal alloys arranged in standalone or tetrode configurations, overcoming the material and geometric limitations of the thermal drawing process. *In vivo* mouse studies validate the P-DOpE probe’s suitability for simultaneous photometry, optogenetics, and electrophysiology over months of chronic recording.

We then used the P-DOpE probe to test whether a local neuronal population regulates its own neuromodulatory tone. The glutamate amplifies noradrenergic effects (GANE) model predicts that an active region of cortex should produce higher noradrenergic signaling, thus creating a local norepinephrine “hot spot”^18^. Our multimodal approach enables us to test this prediction directly, using FP measurement of GRAB_NE_ to monitor NE, optical stimulation of ChrimsonR to depolarize CA1 neurons, and electrophysiology to monitor network LFP and spiking. Our findings support the central hypothesis of GANE: CA1 stimulation accentuates, but does not instantiate, NE release. The P-DOpE probe thus enables low-cost and customizable instrumentation to study how local circuit activity shapes neuromodulatory signaling and behavior.

## Results

### P-DOpE probe fabrication via convergence taper-break

To create the P-DOpE probes with simultaneous photometry, optogenetics, and electrophysiology recording capabilities, we developed a new fabrication approach – convergence taper-break (CTB). Unlike traditional fiber drawing methods in which materials are limited to thermoplastic polymers and low-melting point metals, CTB allows high-melting point materials (such as silica, tungsten, and nichrome wires) to be drawn within a uniform and thin polymer matrix, enabled by the Marangoni effect of viscous materials under a controlled temperature gradient.

Prior to CTB, a cylindrical “mini-preform” (∼1.5–3 mm in diameter, ∼4–8 cm in length) was fabricated from a polycarbonate (PC) rod with multiple hollow channels using an established thermal fiber drawing process (Fig. 1a, b)^17^. Device elements (e.g., silica waveguides, tungsten wires, or nichrome tetrodes) were then inserted into their corresponding channels within the mini-preform (Fig. 1c). The mini-preform step allows for electrode configuration and fiber diameter to be customized with electrode placements optimized for spatial coverage or tetrode configurations.

**Fig. 1.**
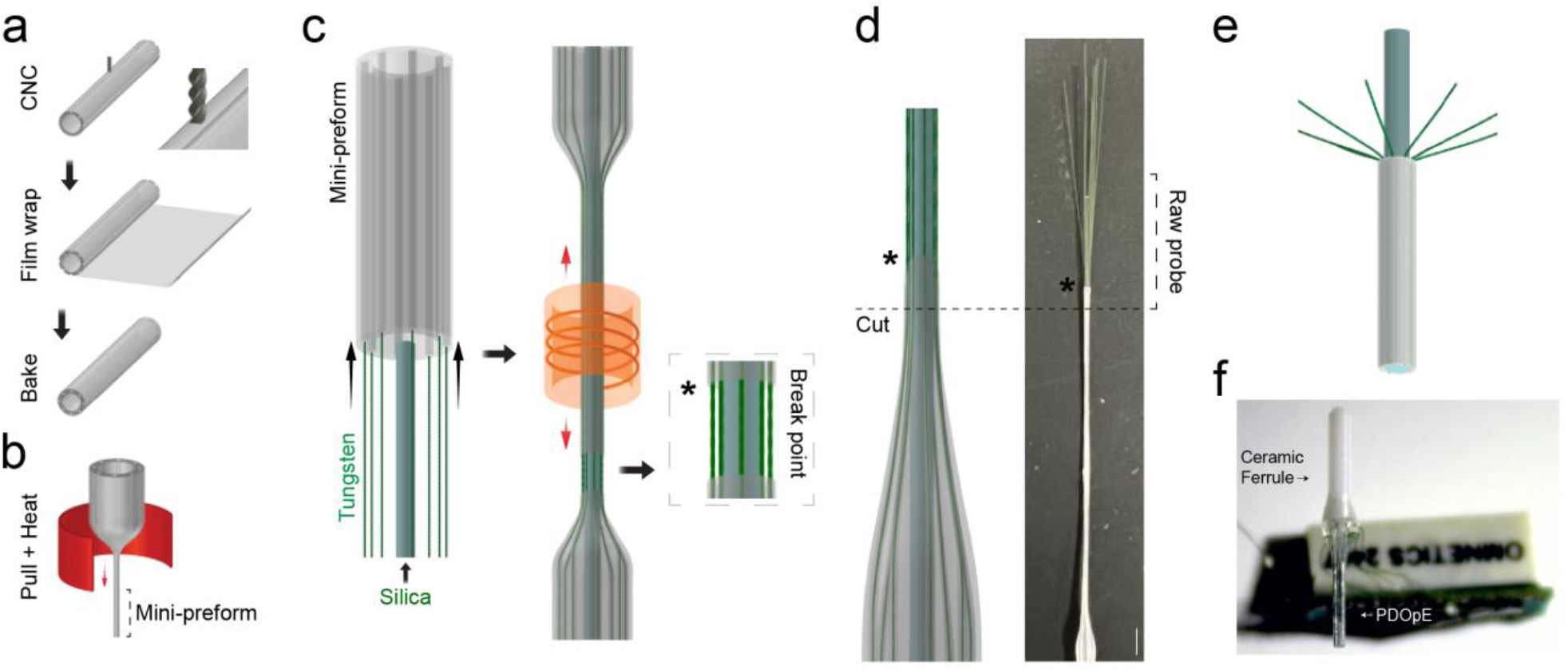
Convergence taper-break (CTB) fabrication of P-DOpE probes. a, Preform fabrication. A polycarbonate (PC) tube is CNC-milled with longitudinal channels (*inset*), wrapped with additional PC film, and thermally consolidated under vacuum (bake) to form the preform. **b**, Mini-preform fabrication. The preform is heated and drawn (pull + heat) in a gradient furnace into a ∼2-mm-diameter mini-preform that preserves the channel layout at reduced scale. **c**, Convergence taper-break (CTB). Device elements — a silica waveguide (*center*) and nichrome electrodes (*outer channels*) — are inserted into the mini-preform, which is then heated by a ring furnace and drawn under tension (*red arrows*) until the thin polymer framework breaks cleanly, leaving the silica waveguide and metal electrodes intact and encapsulated within the polymer. Inset, the break point (*asterisk*). **d**, As-fabricated (“raw”) probe. *Left*, schematic of the drawn structure indicating the cut location (*dashed line*) that separates the sensing tip from the backend; *right*, photograph of the raw probe. Asterisk, break point; scale bar, 5 mm. **e**, Backend geometry. Beyond the break the separated electrodes splay away from the silica/polymer core (asterisk, break point), leaving long lengths of waveguide and wire exposed for connectorization. **f**, Fully assembled ready to implant probe.

During CTB, the assembly was subjected to tensile drawing under a controlled axial thermal gradient established by a ring furnace (1.5 cm in length) surrounding the mini-preform (Fig. 1c). Simultaneous heating and tensile drawing softened the polymer in the necking region, generating a Marangoni stress that overcame capillary instability and produced a uniform device with metal wires positioned precisely around the silica core within a thin polymer matrix. As drawing continued, the mechanical stress exceeded the ultimate tensile strength of the polycarbonate (PC), inducing a clean break in the polymer framework while preserving the integrity of the device elements (Fig. 1c, *Inset*).

The resulting structure was cut in the tapered section (Fig. 1d, *dashed line*) and polished using an optical fiber polisher, yielding a uniformly sized device with separated electrodes and silica waveguide in the backend which can be connectorized according to industry standards (Fig. 1e, f). Fiber configuration customizability enabled by the CTB fabrication process is shown in Fig. 2a, b which shows three example configurations (*left*: nichrome tetrodes with silica waveguides and microfluidic channel; *middle*: tungsten electrodes with 400 µm diameter silica waveguide; *right*: tungsten electrodes with 200 µm diameter silica waveguide). The probe’s light delivery capability was verified using a 0.6 wt% agar tissue phantom (Fig. 2c) and electrode impedance was suitable for single unit recordings (Fig. 2d; channel mean = 140 kΩ at 1 kHz).

**Fig. 2.**
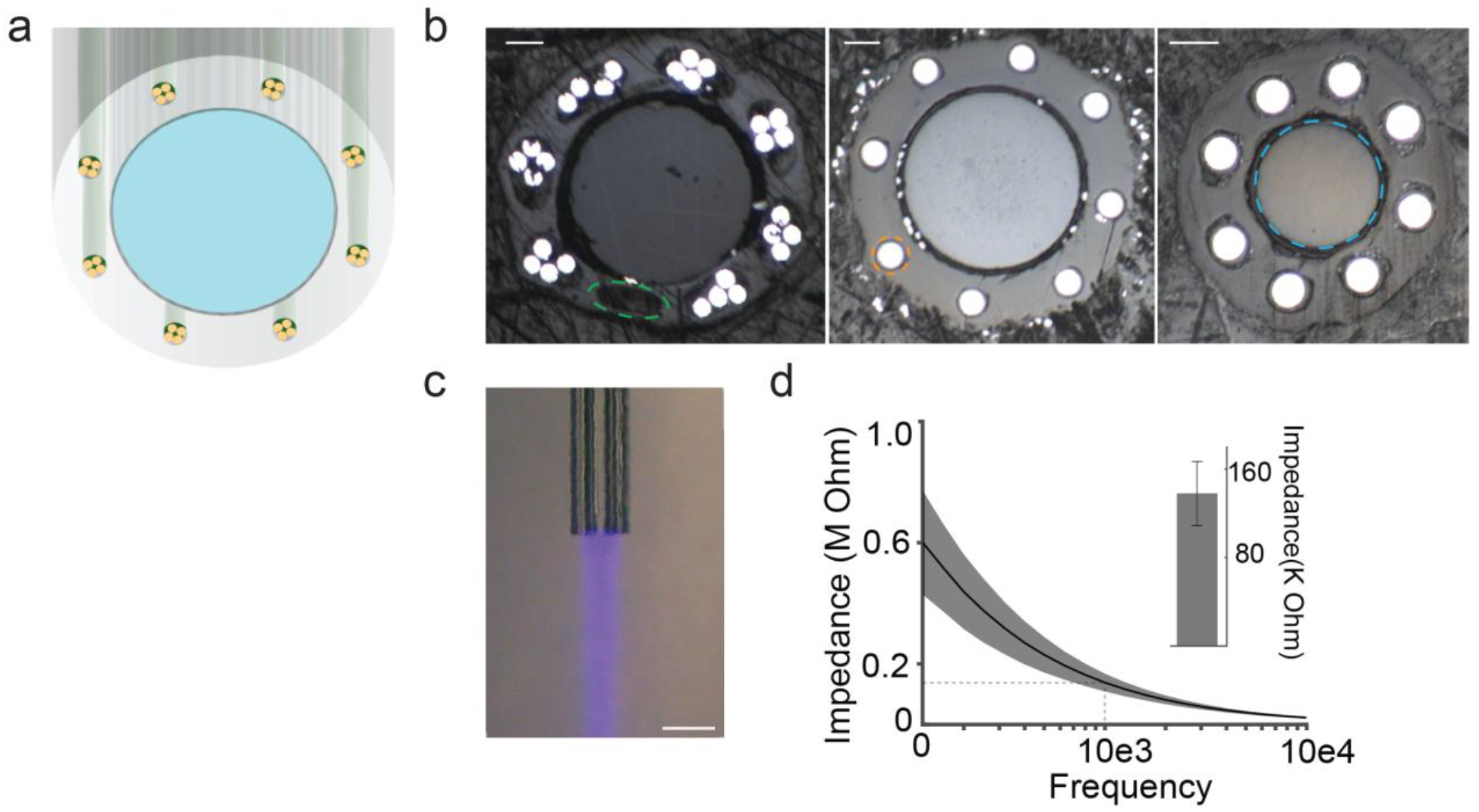
P-DOpE probe configurations and benchtop characterization. **a**, Cross-section of the probe showing the central silica waveguide surrounded by seven nichrome tetrodes (twenty-eight electrodes) within the polymer matrix. **b**, Cross-sections of three example P-DOpE configurations. *left*: seven nichrome tetrodes, a 400-µm silica waveguide, and a microfluidic channel (green); *middle*: eight 50-µm tungsten electrodes (orange) around a 400-µm silica waveguide; *right*: eight 50-µm-diameter tungsten electrodes around a 200-µm silica waveguide (blue). Scale bars, 100 µm. **c**, Light emission through the integrated silica waveguide (0.6 wt% agar tissue phantom), demonstrating optical delivery for stimulation and photometry. Scale bar, 500 µm. **d**, Electrode impedance spectrum of the 50-µm tungsten electrodes (n = 8; line, mean; shading, s.d.). Dashed lines and inset mark the value at 1 kHz (8-channel mean ≈ 140 kΩ).

### In vivo local field potential and single unit responses to optogenetic activation of CA1 with P-DOpE

To assess whether P-DOpE probes are suitable for optogenetic and FP applications, we prepared mice with co-expression of the red-shifted channelrhodopsin ChrimsonR^19^ (AAV5-hSyn-Chrimson) and the NE biosensor GRAB_NE_ (AAV9-hSyn-NE2h)^20^ in dorsal CA1 neurons. In mice chronically implanted with the P-DOpE probe directed at area CA1 (Fig. 3a), we were able to measure multi-unit spikes as well as isolated single units (3-7 units per subject, mean = 5.3 units) during recordings conducted one-week post-implantation (Supplementary Table 1). Spike auto-correlations, cross-correlations, and waveforms suggest most units were not pyramidal cells, which aligns with the more dorsal placement of the probe sensing surface (Fig. 3b). Next, we assessed chronic recording stability by taking measurements across 10 weeks. Despite a reduction in unit yield, the isolation quality of the remaining units improved, as reflected by decreasing L-ratio from Week 1 to Week 11 (Fig. 3c), consistent with resolution of gliosis after initial implantation^21^. Therefore, the P-DOpE probe can capture single and multi-unit responses across months in the awake-behaving subject.

**Fig. 3.**
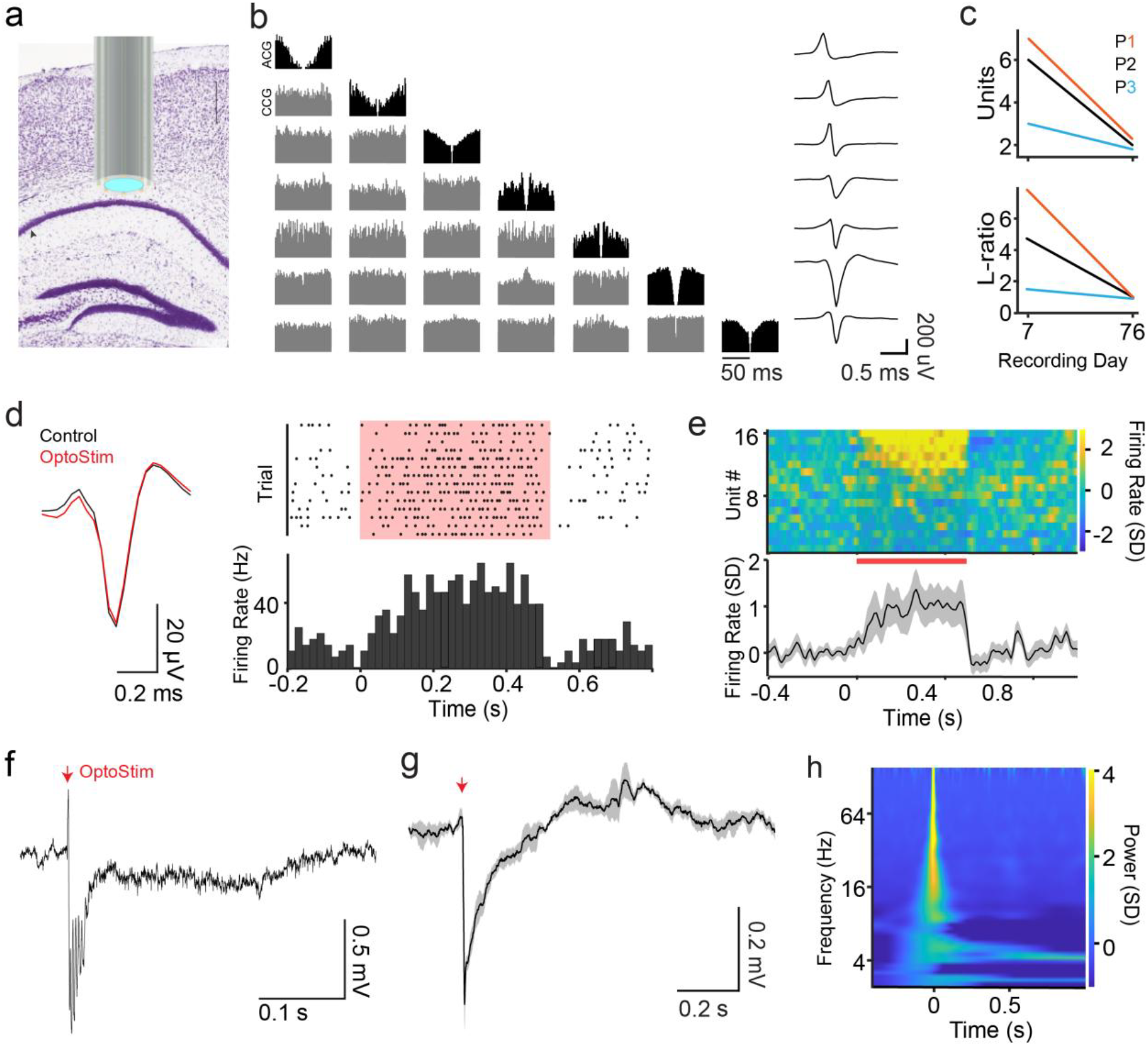
In vivo electrophysiology and optogenetic responses from hippocampal CA1 in chronically implanted, freely moving mice. **a**, Nissl-stained coronal section showing P-DOpE placement just dorsal to the CA1 pyramidal layer (schematic overlay; cyan, light-emitting tip) (*Image from Allen Institute Brain Atlas Creative Commons License*). Arrow indicates pyramidal layer, scale bar 500 µm. **b**, Auto-correlograms (*diagonal*), cross-correlograms (*off-diagonal*), and the mean best-channel waveform for each simultaneously isolated unit. **c**, Chronic recording stability: mean unit count (*top*) and mean L-ratio (*bottom*) per subject on post-implantation Week 1 and Week 11; colored lines, subjects (P1–P3). Lower L-ratio indicates better isolation. **d**, Representative single-unit response to optogenetic stimulation. *Left*, mean spike waveform during (red) and outside (black) stimulation; *right*, raster and peri-event time histogram for the same unit across stimulation trains (shaded). Scale bars, 20 µV and 0.2 ms. **e**, Population response to optogenetic stimulation. *Top*, z-scored firing rate for all units (heat map; red bar, stimulation); *bottom*, mean firing rate ± s.e.m. across units. **f**, Representative wideband extracellular recording (0.1–6,000 Hz) around optogenetic stimulation (*red arrow*). **g**, Mean evoked LFP (± s.e.m.) across trials, baseline-subtracted using the pre-stimulus interval (*red arrow*, stimulation onset). **h**, Wavelet spectrogram of the evoked response, z-scored power relative to the pre-stimulus baseline.

Next, we established the ability of P-DOpE to measure the neuronal response to focal optogenetic stimulation. Optical stimulation of ChrimsonR (638-nm, 500-ms, 60-90 s ISI, output power = 1 mW, corresponding to an irradiance of ∼8 mW/mm^2^ at the 400 µm fiber core) caused a diversity of firing rate responses across the individual units (Fig. 3d, e). Notwithstanding, the trial-averaged population firing rate response revealed a robust, time-locked increase that lasted the duration of the stimulation (Fig. 3e, *bottom*). Evoked LFPs were also observed, characterized by an early rapid negative deflection and high-frequency oscillation followed by a slower return toward baseline and rebound coinciding with stimulation termination (Fig. 3f–h).

### Same-field recordings reveal an optogenetic stimulation artifact independent of NE binding

Having established effective light delivery and electrical recordings with the P-DOpE probe, we next sought to link neuronal excitation with FP measurements of noradrenergic signaling. First, we conducted dose response curves to calibrate network responses across varying light intensities. As expected, higher light power was associated with larger evoked LFP responses (Fig. 4a). In parallel, FP recordings revealed intensity-dependent decreases in fluorescence (Fig. 4b), yielding a strong correlation between the early phase (0 to 25-ms) of the LFP deflection and the early phase of the FP response (0 to 5-s) (*Linear Mixed Effects Model: β = 0*.*0017, t(168) = 13*.*33, p = 4*.*0 × 10*^*-28*^; *n = 3 mice, each Pearson R > 0*.*7*; Fig. 4c). To determine whether the optically induced decrease in fluorescence reflected inhibition of NE release or other physiological changes unrelated to NE binding (e.g. changes in blood oxygen^22^, pH^23^, and/or hemodynamic volume), we injected yohimbine (5 mg/kg), which antagonizes the α_2_-adrenergic receptor, from which the GRAB_NE_ reporter is derived. Isosbestic-corrected GRAB_NE_ emission intensity, referred to as Signal_NE_, was enhanced around the I.P. injection (Fig. 4d),thus serving as a positive control for the non-antagonized sensor, but was completely absent when mice were later transferred to a novel context, a challenge that triggered a phasic increase in Signal_NE_ in the absence of yohimbine (Fig. 4d)^24^. On the other hand, yohimbine had no effect on the fluorescent changes around optogenetic stimulation, strongly suggesting that this stimulus-locked artifact does not reflect NE binding (Fig. 4e).

**Fig. 4.**
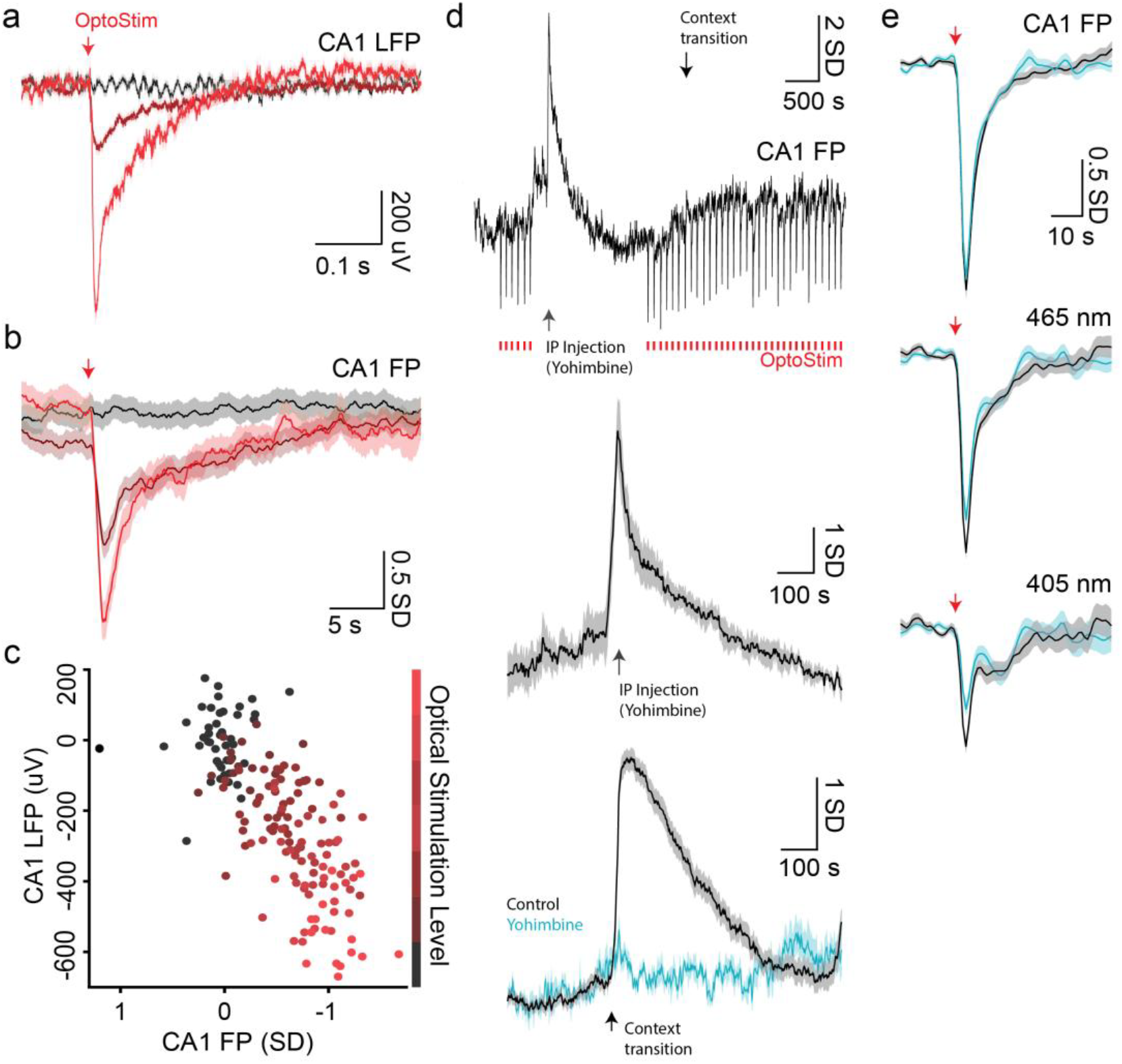
Same-field optogenetic stimulation produces a substrate-independent fluorescence artifact. **a**, Optogenetically evoked CA1 LFPs are intensity-dependent: trial-averaged LFP at each of 7 light intensities (*color gradient*, dark → red with increasing power). *Red arrow*, optogenetic stimulation onset (638-nm, 500-ms). Shaded regions, s.e.m. **b**, Simultaneously acquired GRAB_NE_ fiber photometry (CA1 FP) shows intensity-dependent decreases in fluorescence to the same stimuli (same intensity color scale and stimulation marker as *Panel a*). Shaded regions, s.e.m. **c**, The early evoked LFP (0–25 ms) and the early FP response (0–5 s) are strongly correlated across the intensity series. Each point is one trial, colored by optical-stimulation level (For each mouse, Pearson R > 0.70; 8.1 ± 5.05 trials per intensity per subject, n = 3 mice). **d**, Pharmacological controls in an example session (CA1 FP). *Top*, full session showing optogenetic-stimulation epochs (*red dashes*), I.P. yohimbine injection (5 mg/kg), and transfer to a novel context. *Middle*, Average FP response to injection: yohimbine delivery evokes a clear Signal_NE_ transient, confirming the sensor reports endogenous NE (positive control for the sensor). *Bottom*, expanded view of context transfer: the large context-evoked NE response seen in control sessions (black) is abolished under yohimbine (cyan), confirming that α_2_-receptor blockade eliminates NE detection (positive control for the antagonist). Shaded regions, s.e.m. **e**, The stimulation-evoked fluorescence transient does not reflect NE binding. CA1 FP response to optogenetic stimulation (*top*), and the underlying 465-nm experimental (*middle*) and 405-nm isosbestic (*bottom*) photometry channels, each compared with (cyan) and without (black) yohimbine; red arrow, stimulation onset. The transient is statistically unchanged by yohimbine and is present in the NE-insensitive 405-nm isosbestic channel, therefore it is considered a stimulus-locked artifact. n = 4 mice (all yohimbine sessions).

To correct for this confound, we relied on the fact that the response was highly stable over the hour of recording (time-invariant) and was identical under reporter antagonism. Simple subtraction of the average impulse response function (IRF) measured in the home cage yielded a corrected Signal_NE_ that was free of distortions (Supplementary Fig. 1), suggesting that the stimulus-locked artifact distorted the Signal_NE_ in a linear, time-invariant fashion.

### CA1 stimulation amplifies noradrenergic release

The ability of P-DOpE to deliver light-manipulation to the same neurons under electrophysiological observation enabled calibration of optogenetic intensity across subjects, while the pharmacology allowed correction for evoked fluorescence changes unrelated to substrate binding. With these controls in place, we were able to test the central prediction of the GANE hypothesis^18^: that local activity should modulate ongoing NE release. To evoke endogenous phasic NE release, subjects were transferred to a novel context and local activity was enhanced by optogenetically stimulating CA1 neurons before and during context transition; control sessions involved transfers without stimulation. We found that CA1 stimulation did not cause NE release in the home cage (Fig. 5a). However, stimulation significantly modulated the NE response around context transition, causing a 54.3 ± 7.0% increase in total NE released (AUC; Fig. 5b) after context transition, relative to within-subject controls without light (*t(3) = 10*.*0, p = 0*.*002*). These results support the central prediction of the GANE hypothesis, demonstrating that local hippocampal activity can amplify, but does not independently induce, noradrenergic signaling.

**Fig. 5.**
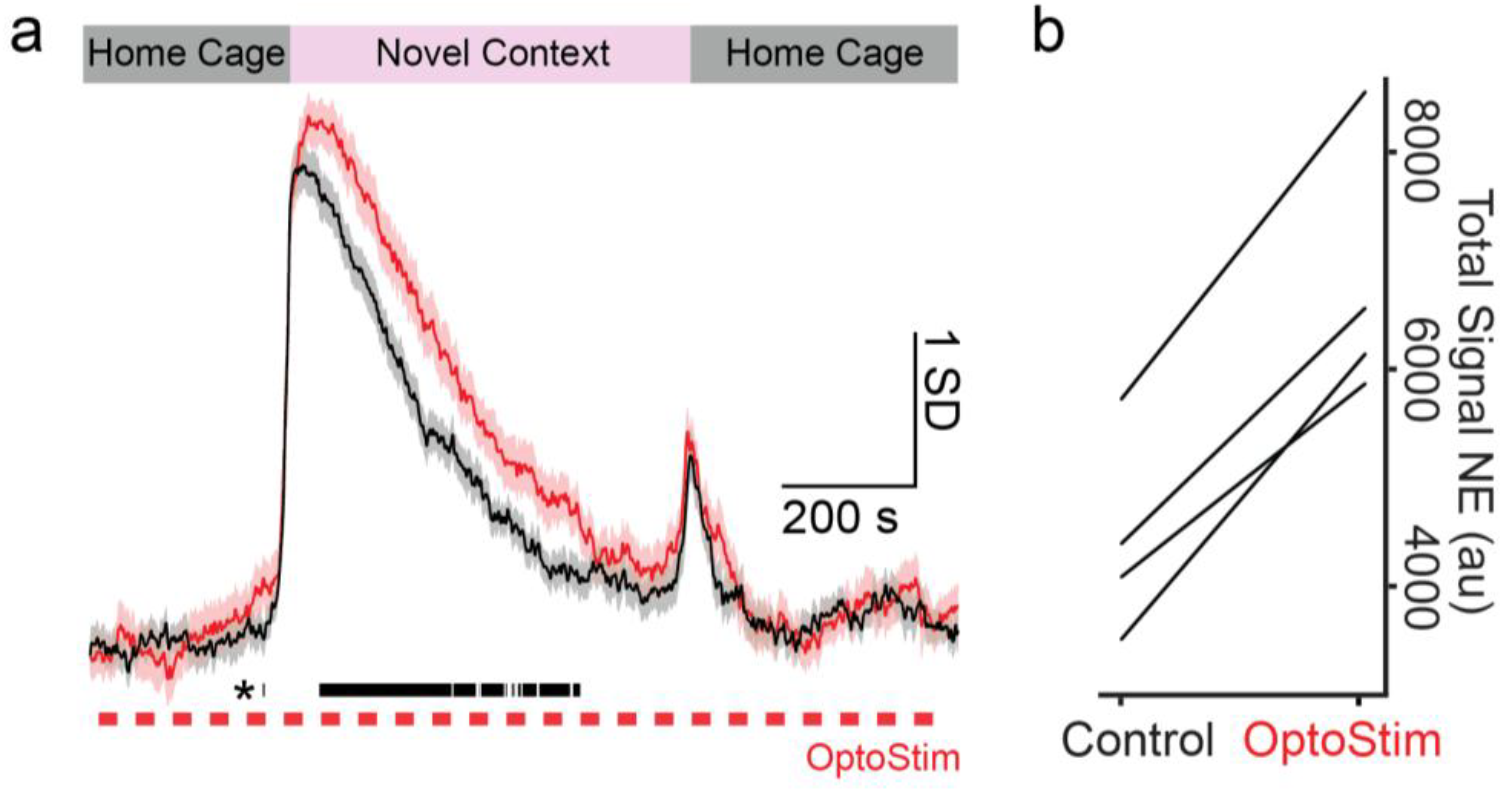
CA1 stimulation amplifies the context-evoked noradrenergic response, supporting the GANE hypothesis. **a**, Signal_NE_ across a home-cage → novel-context → home-cage session. Optogenetic stimulation of CA1 (red; OptoStim, *red dashes*) prolongs and amplifies the context-evoked NE response relative to within-subject control sessions without stimulation (black). Stimulation begins 5 min before context transfer; the conditions do not differ before the transfer-evoked surge. Bar and asterisk, window of significant difference (p< 0.05). Scale bars, 1 SD and 200 s. **b**, Total NE per subject (area under the curve) was greater with than without optogenetic stimulation (each line, one mouse; 54.3 ± 7.0% increase; *t(3) = 10*.*0, p = 0*.*002*).

## Discussion

In this work, we report a new method, CTB, for the fabrication of P-DOpE probes consisting of vastly different materials such as silica and high-melting point metal wires in a thin polymer matrix. In the conventional thermal drawing process, materials should mostly consist of thermoplastic polymer and low-melting point metal or metal alloys with compatible rheological properties to avoid structural defects and fiber breakage during the drawing process. This specific combination ensures the core and cladding flow smoothly together in a laminar fashion, preventing capillary instability. On the other hand, in CTB, high-melting point silica and metal wires are drawn within a thin polymer matrix. This is attributed to the controlled thermal gradient in the necking region, which induces a large Marangoni stress overcoming the capillary instability, thus creating a thin and uniform device with precisely and closely placed device elements. Using the P-DOpE probes, we found that increasing the spiking activity of CA1 neurons amplifies the noradrenergic signaling related to a context change, but does not induce NE release in the absence of a phasic trigger. These results support the predictions of the GANE model^18^ in a preparation where the activity of neurons can be experimentally controlled and monitored alongside local NE levels.

### Comparison with other devices

Previous efforts to integrate photometry, optogenetics, and electrophysiology into a single neural probe have typically relied on implanting separate fiber optic stubs and electrophysiology probes, or bonding them together prior to implantation^9,10^. These approaches are difficult to scale or reproduce, and precise co-localization of optical and electrical recording sites remains a major challenge.

Thermal drawing processes (TDP) provide a low-cost, scalable route to fabricate multifunctional fiber-based neural probes by drawing macroscopic preforms into microscale fibers^11-16^. However, TDP devices face two key limitations: (1) a trade-off between small probe diameters (needed to minimize tissue damage) and larger back-ends required for reliable electrical and optical connections, and (2) material constraints, as the process typically requires polymers and low-melting point metals (e.g., BiSn, Sn) with matched viscosities, excluding high-performance materials like silica, tungsten, or nichrome wires due to their high processing temperature.

Thermal tapering (TTP) addresses the geometric constraint by introducing an intermediate “mini-preform” (∼2–3 mm diameter) that is later tapered into a probe with a small sensing tip and a larger backend for connectorization^17^. However, TTP does not resolve the material incompatibility. While convergence fiber drawing has been proposed to integrate dissimilar materials^14,25^, co-drawing silica with multiple metal electrodes at small scales has been hindered by polymer instability during the thermal drawing process.

CTB resolves this by controlling the axial thermal gradient to ensure even and precise polymer encapsulation of the silica waveguide and metal wires. Marangoni instability arises when temperature gradients induce surface tension variations, driving fluid flow from hotter (lower surface tension) to colder (higher surface tension) regions. During the CTB, the Marangoni stress overcomes the capillary instability, minimizing the breakage and discontinuities in the viscous thin polymer layer. Therefore, uniform devices with high-melting point metal wires precisely and closely placed around the silica core within a polymer matrix can be fabricated for the first time. The process also significantly enhances device modularity, allowing the same mini-preform configuration to be used for multiple different device designs by simply varying the inserted sensing elements.

A limitation of the device is that the electrode contacts are necessarily flush with the optic waveguide. In the present experiments, the P-DOpE probe was implanted just dorsal to the CA1 pyramidal cell layer, so as to minimize damage to neuronal cell bodies. Such placement limits how action potentials may be captured. Indeed, a subset of isolated units displayed positive-going (non-negative) extracellular waveforms (Fig. 3b). Rather than reflecting poor isolation, such non-negative waveforms are increasingly understood to arise from non-somatic compartments^26-28^. The polarity of these units is therefore consistent with an axonal, dendritic, or return-current origin, as documented across recording technologies including Neuropixels recordings from humans^29^ and ultra-high-density silicon probes^30^.

### Potential sources and roles of GANE

While prior findings are consistent with local control of noradrenergic tone, to our knowledge, this is the first *in vivo* demonstration that cortical activity within a healthy physiological range can bias local NE signaling. Prior studies, including those in the awake rat, found that induction of LTP through electrical stimulation of the perforant path causes local and sustained changes in hippocampal NE release^31,32^. The use of electrical stimulation in these studies confounds whether the NE release was due to direct stimulation of LC terminals versus local control of NE release by hippocampal principal neurons, *per se*. In our experiments, it is unlikely that LC was inadvertently stimulated as we did not see any retrograde viral expression in the LC, nor were we able to evoke phasic NE release upon a tonic background. Others have shown that glutamatergic activation of hippocampal slices causes NE release^33^ which can also be observed in a reduced hippocampal synaptosome preparation^33,34^, thus indicating that glutamate can induce NE release from LC axonal boutons. Therefore, one potential mechanism for GANE is glutamate enhancing NE release probability. An alternative possibility is that hippocampal activity may slow NE clearance, as the norepinephrine transporter (NET) has been shown to be less effective when cells are depolarized^35^. Brief stimulation of the LC causes sustained NE release in the hippocampus^36^ and prefrontal cortex^37^, but not LC^36^, potentially due to regional variations in NET expression. Finally, polysynaptic drive of LC somata is possible. Induction of hippocampal seizures causes hippocampal NE release and modulation of LC unit activity^38^, suggesting that at least under these extreme levels of activity, such cross-regional synchronization occurs.

Our previous work showed that the decay of Signal_NE_ after context change was well approximated by a decaying exponential^24^, suggestive of first-order pharmacokinetic clearance dynamics dictated by the efficacy of NET. Importantly, learning, which is associated with decreased activity of glutamatergic neurons over days^39^, accelerated the SignalNE decay rate, potentially due to the removal of the GANE phenomenon observed here. Future studies, ideally in which NE is sampled from multiple locations, will be needed to elucidate the anatomical specificity and underlying mechanism driving the observed amplification of noradrenergic signaling.

CA1 activation was insufficient to elicit NE release in the home cage, yet robustly amplified NE signaling once the phasic trigger of context transfer was present. This pattern suggests that hippocampal activity is not a primary driver of noradrenergic output and that local circuit activity can bias neuromodulatory signaling only when the LC–NE system is already engaged, consistent with models in which neuromodulators integrate local salience with global brain state^18^. These results support a role for hippocampal activity in shaping the neuromodulatory signal, rather than generating a nonspecific arousal signal, and align with theoretical accounts in which NE acts to selectively amplify encoding and consolidation processes triggered by salient events^40-48^.

The combined functionality of the P-DOpE probe enabled testing of key predictions of the longstanding GANE hypothesis, demonstrating its utility for optogenetic manipulation and multimodal readout within a single low-cost, customizable device.

## Methods

### Preform fabrication

Preform fabrication began with CNC milling eight 3.6 x 3.6 mm rectangular channels on a polycarbonate (PC) tube (OD: 26 mm; ID 19 mm) (McMaster-Carr, IL) which was then wrapped with PC films to a final diameter of 28.6 mm and consolidated at 190° C in a vacuum furnace. A 19 mm diameter Teflon rod (McMaster-Carr, IL) was inserted in the center hole of the preform to maintain preform geometry during consolidation. After that, the Teflon rod was removed leaving a hollow channel in the center.

### Mini-preform fabrication

The preform was mounted in a 3-temperature-zone (top, middle, bottom) gradient furnace. The zones’ temperatures were set at 150 °C, 275 °C and 120 °C respectively (Note: these are readouts of the thermocouples mounted in the furnace and thus may not accurately represent the actual temperatures). The preform was then heated until softened and thermally drawn into 2 mm diameter mini-preforms via a capstan motor.

### Convergence taper-break

The sensor elements were inserted into their corresponding empty channels on the mini-preform with the wires (Tungsten wires were obtained from California Fine Wire Co, CA; nichrome wires were obtained from AM-Systems Inc, Sequim, WA) in the 8 outer channels, and the silica waveguide (Thorlabs, NJ) in the center. The mini-preform was then loaded into a custom-built thermal tapering setup consisting of optomechanical components, heated to 230° C to soften the mini-preform, and pulled until the softened polymer runs out and shears, leaving long sections of wire and waveguide exposed for ease of connection. The probe is then cut, and the probe tip polished down to 1 μm roughness via an optical fiber polisher (KrellTech, NJ).

### Probe assembly

To fully assemble the probe, short sections of the tungsten wires insulation layers were removed prior to the connectorization to ensure electrical contact with the solder. The silica waveguide was fitted within a commercial 1.25 mm diameter ceramic ferrule for coupling to standard optical components. The wires were soldered onto the Printed Circuit Board (PCB), and a 1.5 mm diameter ferrule (Thorlabs, NJ) was glued onto the optical fiber. The ferrule and optical fiber tip were then polished down to 1 μm roughness via an optical fiber polisher (KrellTech, NJ). Electrode impedance was measured via Electrochemical Impedance Spectroscopy (EIS) in saline using a potentiostat.

### In vivo mouse experiments

C57BL/6J mice (n = 4 male littermates) were implanted at 2-3 months old. Data were acquired 2-12 weeks after viral injection. Mice underwent two surgeries, the first to deliver the GRAB_NE_ sensor^20^ and ChrimsonR opsin^19^ via infusion of an AAV cocktail, and the second to implant the P-DOpE probe. After viral injection, animals were housed individually on a regular 12:12-hr light:dark schedule and tested during the light cycle. Following one week of recovery from the second surgery, mice were recorded at most 5 days/week before being euthanized with a sodium pentobarbital cocktail (FatalPlus®, 300 mg/kg I.P.) and transcardially perfused with 4% paraformaldehyde. All experimental procedures were performed in accordance with the National Institutes of Health Guide for Care and Use of Laboratory Animals and were approved by the University of New Mexico Health Sciences Center Institutional Animal Care and Use Committee.

### Viral injections and fiber implantation

Mice were deeply anesthetized with isoflurane (1.5-2% in pure oxygen) and GRAB_NE_ (AAV9-hSyn-NE2h; titer: ≥ 5×10^12^ vg/mL, WZ Biosciences, MD USA)^20^ and ChrimsonR (AAV5-hSyn-ChrimsonR-tdT; titer: ≥ 7×10^12^ vg/mL, Addgene #59171)^19^ were delivered by injecting unilaterally into the left dorsal hippocampus (A/P: -2.3, M/L: -2.0, D/V: -1.4 from the brain surface). The AAV cocktail was injected at a volume of 150-nL and a rate of 30 nL/min using a Nanoliter 2020 Injector (WPI, FL). At least 1 week later, the P-DOpE probe was implanted dorsal to the injection site. To secure the probe to the subject, the surface of the exposed skull was covered with C&B Metabond® (Parkell, NY), and the sides of the exposed fiber-optic cannula were coated in Unifast LC dental acrylic (SourceOne Dental, Inc, AZ) for stability. Finally, clips (Neuralynx, AZ) were added to minimize motion artifact due to slippage at the mating sleeve. Postoperatively, animals received a single injection of 0.1-mg/kg of buprenorphine (S.C.) and again as needed for the next 1-3 days.

### Fiber photometry recording, electrophysiology, and optogenetic stimulation procedures

Prior to the first recording session, we allowed a minimum of 2 weeks from the viral injection procedure to allow the virus sufficient time to transfect and express. Signals were captured with a LUX RZ10X processor running the Synapse software (Tucker-Davis Technologies, FL). Experimental (465 nm, carrier frequency = 330 Hz) and isosbestic (405 nm, carrier frequency = 210 Hz) wavelengths were combined using a fluorescent MiniCube (FMC5_IE(400-410)_E(460-490)_F(500-540)_O(580-680)_S; Doric, QC Canada) and delivered to the subject with a 4-m low autofluorescence mono fiber-optic patch cord (core = 400-µm; NA = 0.57; Doric, QC Canada). Excitation intensity of the isosbestic and experimental wavelengths was adjusted to equalize emission intensity, which was sampled at 1017.3 Hz.

Neural signals were captured at 30-kHz on a 32-channel Intan headstage connected to the RHD Evaluation Board (Intan Technologies, CA).

For optogenetic stimulation, red light (638-nm; Thorlabs L638P150) was delivered via the patch cord. Light pulses (1-mW) were given for 500-ms either at 60-s±6-s or 90-s±9-s (longer durations were used to isolate the stimulation related artifact).

### Behavioral procedures

To measure endogenous NE release, mice were transferred to a novel context. In each recording session, the following protocol was performed: first, a 10-minute home cage (HC) baseline was captured, then mice were manually transferred to a novel arena (Context A) and back to their home cage for 10 minutes. This procedure was performed again for novel Context B. Different arenas were used for each session which varied in shape, size, and color. Light was either given before and throughout Context A (n = 12 sessions), Context B (n = 13 sessions), or neither (n = 4 sessions).

To control for changes in Signal_NE_ not related to NE, yohimbine (5 mg/kg) was injected (I.P.) to eliminate emission fluctuations related to NE binding^20,24^ (note that higher doses elicited seizures when combined with hippocampal stimulation). Drug sessions had the following structure: 10 minute baseline, 10 minutes of optogenetic stimulation (90-s ± 9-s ISI), 5-minute baseline, I.P. yohimbine injection, 25-minute baseline, 5-minute stimulation in home cage, and finally context transfer (with concurrent stimulation).

### Spike sorting and post statistical analysis

Single-unit activity was extracted from broadband recordings (30 kHz), and spike sorting was performed using Kilosort1^49^ and subsequently curated manually in Phy2. Spike isolation quality was assessed using L-ratio^50^, isolation distance^50^, and inter-spike interval (ISI) violation rate for each unit. All metrics were derived from a global principal component analysis (PCA) of raw spike waveforms extracted across all electrode channels, retaining the top 12 principal components to enable valid Mahalanobis distance computation across clusters. Chronic recording stability of the single unit was assessed by computing the mean L-ratio per unit per session and the mean unit yield per probe across the implantation period. Two time windows were defined for comparison: an early period (Week 1, Day 7 post-implantation) and a late period (Week 11, Days 77–86 post-implantation).

### Local field potential (LFP) analysis

LFP data were downsampled from 30 kHz to 1250 Hz. The channel with the strongest evoked response was selected for analysis. Each trial was baseline-corrected by subtracting the mean signal amplitude during the pre-stimulus window (−400 to 0 ms). Trials were averaged first within each animal across sessions, and the grand mean and standard error were computed across animals (n = 4 mice).

Time-frequency analysis was performed on each animal’s mean LFP trace using a continuous wavelet transform (CWT) with a Morlet wavelet (ω_0_ = 6), spanning 2–150 Hz. Instantaneous power was computed as the squared modulus of the complex wavelet coefficients. Power at each frequency was normalized to a z-score relative to each animal’s pre-stimulus baseline (mean and standard deviation computed across −400 to 0 ms). The resulting z-scored spectrograms were then averaged across animals (n = 4 mice).

### Estimation of Signal_NE_

The demuxed experimental and isosbestic signals both exhibited evidence of photobleaching, though with different decay rates. Therefore, we fit a double exponential to the first 10 minutes of each signal to estimate and extrapolate a mean signal which was subtracted from the observed emission intensities. Next, the isosbestic was scaled to the experimental signal using standard linear regression. The isosbestic was then subtracted from the experimental signal, and the mean and standard deviation were calculated over the first 10 minutes. These values were used to normalize Signal_NE_ which is measured in terms of baseline standard deviations from the baseline mean. Finally, the signal was smoothed with a Gaussian kernel (1-s standard deviation).

### Stimulus locked artifact correction

Since the home-cage response of GRAB_NE_ to optogenetic stimulation was equivalent with and without yohimbine, we inferred that these changes in emission reflected a stimulus-locked artifact impulse response function (IRF). This response was stable across the 1 hr of recording. Linear subtraction of the home-cage-derived stimulus-locked artifact IRF yielded smoothly decaying Signal_NE_ during context transfer, indicating that the stimulus-locked artifact distorted Signal_NE_ in a linear, time-invariant manner.

### Data and code availability

All analyses were conducted using MATLAB versions R2018a and higher with the following add-ons: Statistical Toolbox, CellExplorer (https://cellexplorer.org/)^51^, McKenzieNeuroLab Toolbox (https://github.com/McKenzieNeuro/McKenzieLab/tree/main/FiberPhotometry), and Buzcode Toolbox (https://github.com/buzsakilab/buzcode).

## Supplementary information

**Supplementary Fig 1.**
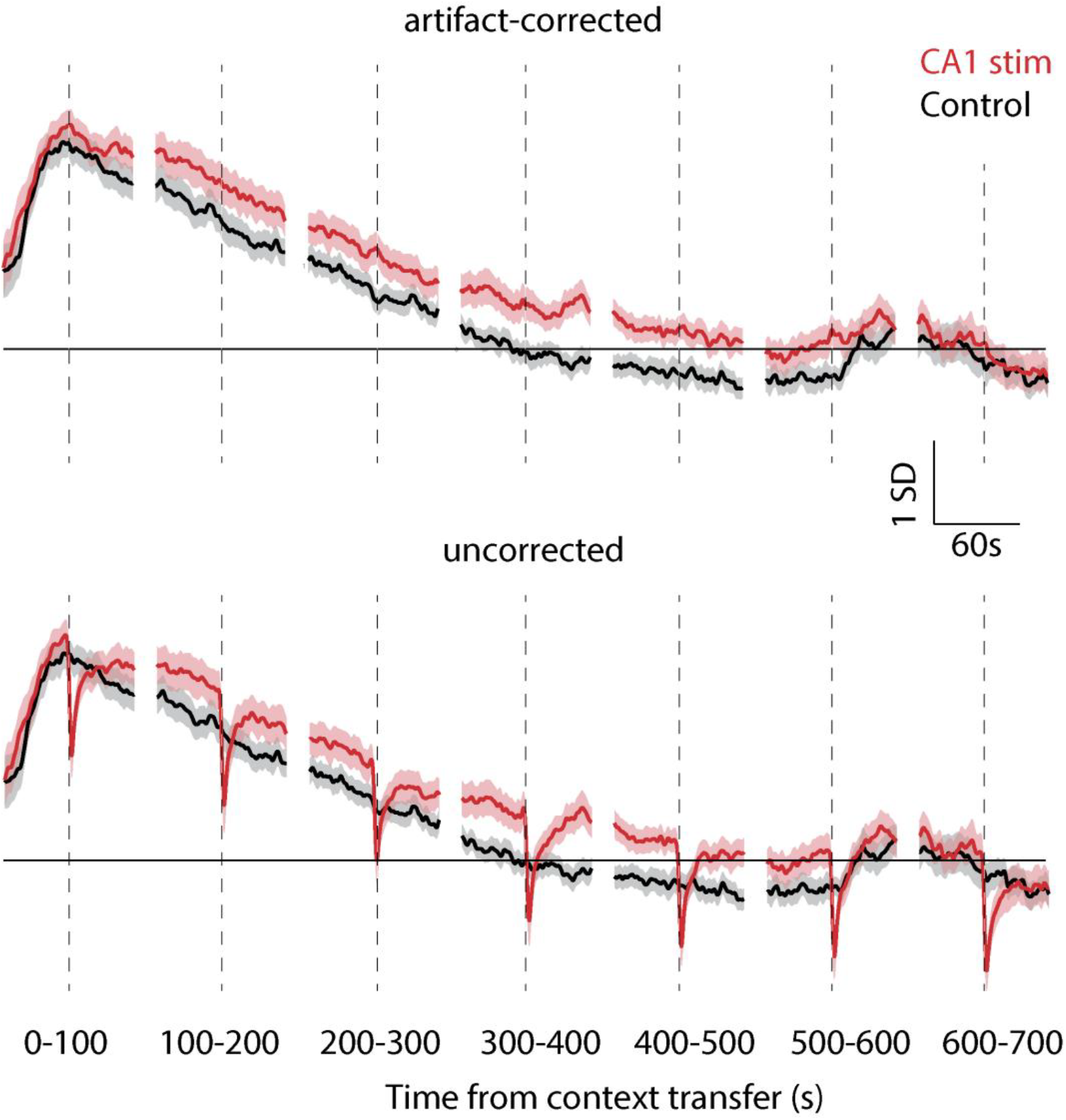
Correction of the context-evoked Signal_NE_ response for the home-cage stimulus-locked artifact. Signal_NE_ binned in 100-s windows from context transfer (0–700 s) for sessions with CA1 optogenetic stimulation (red) and within-subject control sessions without stimulation (black); shaded regions,s.e.m. *Top*, after subtraction of the home-cage stimulus-locked artifact impulse response function (IRF); bottom, the same data without correction. The distortion in Signal_NE_ is visible at each dashed line, where red light was pulsed after context transfer; subtracting the home-cage IRF eliminated the artifact while preserving the stimulation-versus-control difference, indicating that it is linear and time-invariant. Scale bars, 1 S.D. and 60 s.

**Supplementary Table 1.**
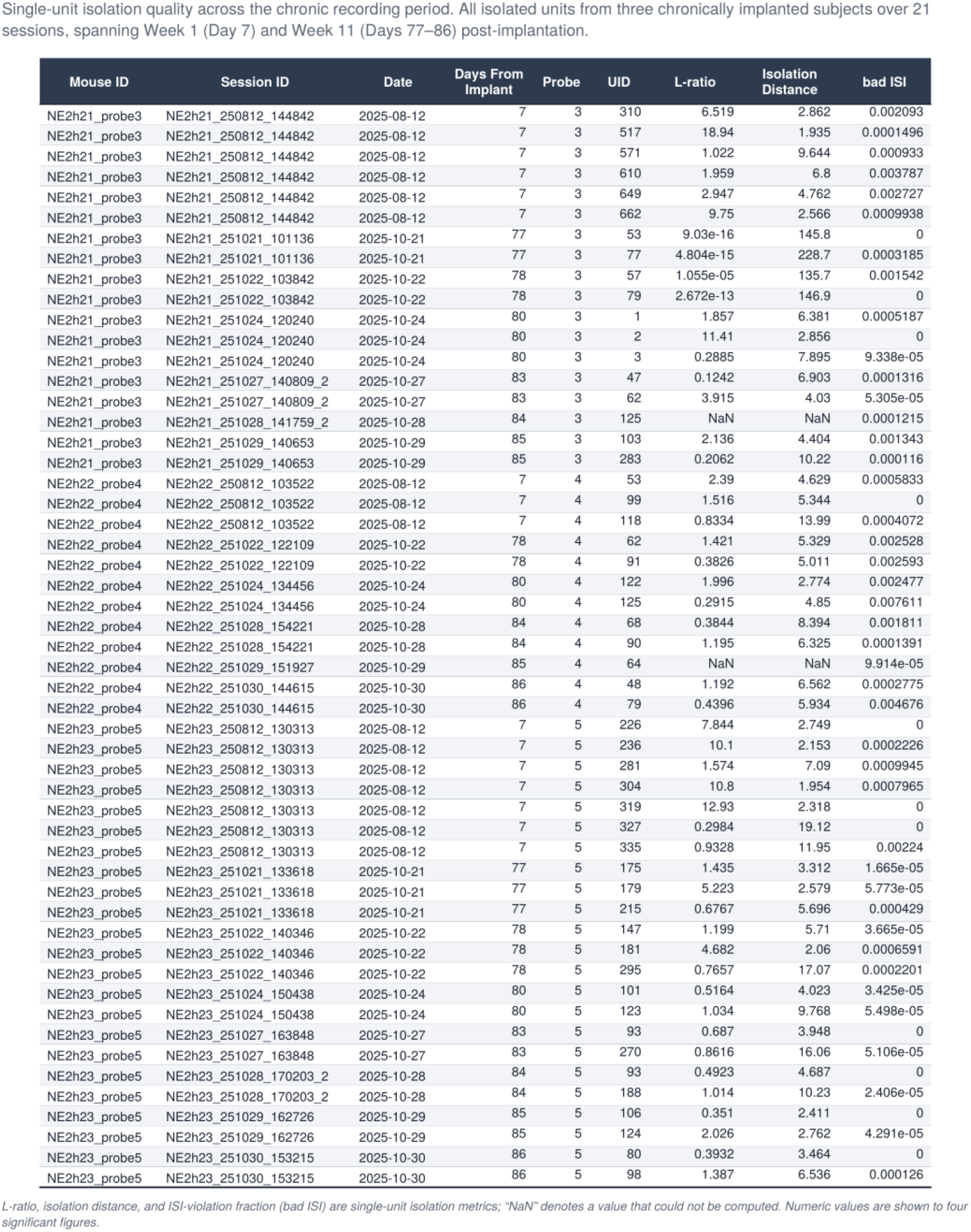
Single-unit isolation quality across chronic recordings. Per-unit cluster-quality metrics for all isolated single units contributing to the chronic-stability analysis in Fig. 3c, listed by subject with units (MouseID; NE2h21–NE2h23), session, recording date, and days post-implantation (Week 1 through Week 11). For each unit (UID) on each probe, L-ratio and isolation distance quantify cluster separation (lower L-ratio and higher isolation distance indicate cleaner isolation), and the inter-spike-interval violation fraction (badISI) reflects refractory-period contamination. Metrics were derived from a global principal-component analysis of spike waveforms across all electrode channels (Kilosort1, curated in Phy2).

## Notes

### Competing Interest Statement

The authors have declared no competing interest.

### Funding

- National Institute of Mental Health: R01MH140076

